# Genome-wide Identification and Expression Profile Analysis of Laccase Family Genes in the *Hypsizygus marmoreus*

**DOI:** 10.1101/2022.12.01.518721

**Authors:** Gang Wang, Cheng Wang, HongBo Wang, Ying Zhu, Yuanyuan Wang, Yu chen, Lin Ma, Zijun Sun, Bobin Liu, Fang Liu

## Abstract

Laccase exists widely in plants and fungi. It is a copper-containing polyphenol oxidase that can degrade lignin, oxidate, and phenolic substances, inhibit heterophytes, promote fruiting body formation, and improve the quality of mushrooms. In this study, 18 laccase genes were identified from the whole genome of a white strain (HM62) of *Hypsizygus marmoreus*, and the mapping, structure, and evolution of laccase genes were analyzed at the whole genome level, while the spatiotemporal expression was evaluated at different developmental stages. The laccase genes mainly distributed on chromosomes 1, 2, 3, 4, 6, 9, and 10, and 9 genes were clustered linearly on chromosome 6, indicating gene doubling. Phylogenetic tree analysis showed that the laccase gene family was divided into three subfamilies. The spatiotemporal expression analysis of the laccase gene family showed that *HmLac09* and *HmLac10* were highly expressed in different periods and might be involved in lignin degradation and fruit body formation, respectively. The expression levels of *HmLac02, HmLac05, HmLac08*, and *HmLac17* genes in gray or gray and white heterozygous strains were higher than those in white strains, which might be related to the difference in lignin decomposition in gray strains, and one of the factors leading to different growth rates. The present study investigated the characterization of the *H. marmoreus* laccase gene family, extending our understanding of laccase mediated fruiting body development and growth rate mechanisms in this fungi.

## 1. Introduction

Laccase is a polyphenol oxidase with copper ions. It is involved in the degradation of lignin together with lignin peroxidase (Giardina et al. 2010), manganese peroxidase, and multifunctional peroxidase and is widely present in plants, insects, fungi, and bacteria (Buddolla et al. 2014). The laccase molecule is composed of a single polypeptide, a copper ion active center, and a sugar ligand. Those from different sources vary in degrees of glycation that use the unique redox ability of copper ions to carry out one-electron oxidation of reducing substrates and reducing oxygen to water (Li et al. 2000). According to the nature of magnetism and spectroscopy, laccase is composed of three conservative copper ion structural domains. The active center of the copper ions is divided into three categories: type I (T1) copper ion (T1-Cu) or blue type copper (T2), type II copper ion (T2-Cu) or type copper, and two type III (T3) copper ions (T3-Cu) or coupling double karyotypes of copper (Li 2014; Hoegger et al. 2006). However, all laccase structures do contain all three types of copper ions. Typically, copper ion has four non-vacancy conserved motifs (L1–L4), which are the marker sequences of laccase from another polyphenol oxidase. These sequences include 10 histidines and 1 cysteine. These amino acid residues combine with the three copper ions of laccase to form ligands that effectuate the physiological roles of laccase (Gold and Alic 1993; Jia et al. 2019; Liao 2017). To adapt to different growth environments, laccase needs various functions; thus, they gradually differentiate into varied homologous genes with different functions (combined sequence and structure analysis of the fungal family). In the evolution of laccase protein, some related functional amino acid residues rarely mutated and became a conserved part, which was used as the identification tag of the gene (the structure and function of fungal laccases).

In addition, laccase genes were involved in lignin degradation, vegetative growth, fruiting body formation, and pigmentation during the growth of edible fungi (Lundell et al. 2010). Laccase gene families have been reported in edible fungi, such as *Coprinopsis cinerea, Auricularia auricula, Flammulina velutipes, Volvariella volvacea*, and *Pleurotus ostreatus*. To date, 17 gene families in *Coprinopsis cinerea*, the largest basidiomycetes laccase gene family, have been identified (Kilaru et al. 2006; Yang 2014; Jiao et al. 2018; Wang et al. 2015; Lu et al. 2015). Several studies have assessed the molecular and functional aspects of this family. The laccase gene of *H. marmoreus* (lcc1), 2336 bp in length containing 13 introns and 14 exons, was cloned, and the phylogenetic tree showed homology with the laccase gene of *Flammulina velutipes*. Interestingly, the laccase activity of the recombinant strain was higher than that of the control, the growth rate of mycelia was significantly increased, the primordia formation was 3–5 days early, and the fruiting body maturity was 5 days higher, indicating that the laccase gene could promote the growth of mycelia and the development of the fruiting body (Zhang et al. 2015). A previous study showed that the activities of laccase and *β*-glucosidase in the primordia stage were significantly higher than those in other states, which might be related to the formation of primordia and promote the early transfer reproductive growth of *H. marmoreus* (Song et al. 2018). Kojic acid is an inhibitor of laccase; a study showed that laccase activity was downregulated by kojic acid during mycelia recovery and color transformation but significantly upregulated during the primary stage, further indicating that laccase is closely related to the fruiting body development of *H. marmoreus* (Zhang et al. 2018).

Hitherto, only a few studies have evaluated the laccase genes due to the lack of a laccase genome and systematic identification, induction, and functional analysis of the laccase gene family. Therefore, in the present study, (1) the laccase gene family was systematically identified, and its structure and chromosomal location were analyzed at the chromosome level of the white strain genome; (2) intraspecific and interspecific evolution of the laccase gene family in *H. marmoreus* was assessed; (3) the spatiotemporal expression of the laccase gene in different tissues and mycelia of *H. marmoreus* in different periods was analyzed.

## 2. Materials and Methods

### 2.1 The materials

Three transcriptomic experiments were carried out to analyze the spatiotemporal expression of laccase genes. In the first experiment, three stages of mycelia post-ripening stage (opening and tiling bacteria), a color turning stage (gray strain turning color, white strain not turning color), and primordia formation were selected during the growth of the *H. marmoreus* grey strain (*Hm61*) and white strain (*Hm88*). (2) In the second experiment, the lid epidermal tissue samples were taken from gray (*Hm61*), white (*Hm88*), and their hybrid progeny (*HMZ5*) strains. (3) In the third experiment, mononuclear mycelium *Hm61_G6, Hm88_W2*, and their hybrid *HMZ5* mycelium were respectively taken. There were three biological replicates per sample in each experiment. Cultivation bag matrix (mass ratio): wood 78%, bran 21%, lime powder 0.5%, gypsum powder 0.5%. Water content 65%.

### 2.2 Sequence retrieval

The white strain HM62-W was the reference genome, which was completed by Genome and Biotechnology Research Center of Strait Joint Research Institute of Fujian Agriculture and Forestry University (NCBI Accession no. JABWDO000000000). The reference genome of the grey strain Haemi51987-8 was published in 2018 (Min et al. 2018). Combined with Hi-C sequencing data, 278 overlapping groups of Haemi51987-8 were interrupted for remounting and gene prediction, resulting in a high-quality genome map named *HM01_Gray. Macrolepiota albuminosa* data from Ensembl Fungi (http://fungi.ensembl.org/index.html) database, while the data of *Wolfiporia extensa* and *Fistulina hepatica* were obtained from NCBI database.

### 2.3 Identification of members of Laccase gene family of *H. marmoreus*

The identification procedures of laccase gene family members of Laccase of *H. marmoreus* were as follows: (1) Using NCBI GenBank (LCC1-LCC17: Bk004111-bk004127), and the amino acid residues of 17 non-allelic laccase genes from *Coprinopsis cinerea* were used as seed sequences. With the help of local Blast software, the sequences with E value less than or equal to 1e^-10^ were used as candidate bases (Hoegger et al. 2006; Camacho et al. 2009). (2) Candidate sequences were compared back to Swissport database, and the sequence with the highest consistency was reserved laccase gene; (3) Through multiple sequence alignment, the genes without laccase marker sequences were deleted; (4) Used the Batch CD-search (https://www.ncbi.nlm.nih.gov/Structure/bwrpsb/bwrpsb.cgi) function of CDD database to predict the domain of candidate genes, and deleted the genes with iron oxidase domain. To avoid the loss of possible laccase gene family members due to incomplete domains, the SMART database (http://smart.embl-heidelberg.de/) was used to verify the existence of three conserved domains. The candidate genes with laccase conserved domain were selected as members of the HmLacs family. The identification method of laccase gene family members of 54 strain of *H. marmoreus, Macrolepiota albuminosa, Wolfiporia Extensa, Fistulina hepatica*, and other edible fungi was the same as above.

Use SignalP 5.0 Server (http://www.cbs.dtu.dk/services/SignalP/) and SecretomeP 2.0 Server (http://www.cbs.dtu.dk/services/SecretomeP/) to predict HmLacs family typical signal peptide and atypical signal peptide.

### 2.4 Gene structure and conserved motif analysis of *HmLacs*

Use of MEME website (http://meme.sdsc.edu/meme/intro.html) tool for *HmLacs* family Motif (Motif) identification and analysis of the Motif width is set to 6-200 residue, the biggest base sequence number for 25, repeat any number of times Using Python scripts to obtain chromosome positions and exon-intron numbers using TBtools for phylogenetic tree gene structure Conservative Motif distribution visualization (Chen et al. 2018).

### 2.5 Chromosomal localization and collinearity analysis of all *HmLacs*

MCScanX software was used to analyze the collinearity and gene duplication events. The E value of blast was less than or equal to 1e^-10^, and the other parameters were default parameters (Wang et al. 2012). For tandem repetition file output by MCScanX, further manual identification was carried out, and the identification criteria were as follows : (1) the ratio of shorter sequence length to longer sequence length was large At 70%; (2) The similarity of the two amino acid sequences is more than 70%; (3) The two genes were in 100 KB fragment (Gu et al. 2002); In addition, GGgenes’ R package was used to carry out microscopic collinearity visualization Bedtools with 80 KB as a unit to count the density of genes on chromosomes, and Tbtools was used to display chromosome location (Hall 2010).

### 2.6 Multiple sequence alignment, phylogenetic analysis, and classification of *H. marmoreus* laccases

The intraspecific (white and gray strain) and interspecific (*Coprinus cinereus, Pleurotus ostreatus, Flammulina velutipes, Lentinula edodes, Volvariella volvacea, collybia albuminosa, Wolfiporia Extensa, Fistulina hepatica*) laccase gene family phylogenetic trees were constructed respectively, and the protein sequences of laccase gene in *Arabidopsis thaliana* were outgroup. Phylogenetic tree construction was carried out with the help of the finsuite manager plug-in (Zhang et al. 2020): (1) multi-sequence alignment of protein sequences using the normal alignment mode of MATTF; (2) deletion of vacancies using trimAl and retention of conserved amino acid residues; (3) ModelFinder Software selects the best protein evolution model (Lanfear et al. 2017) (4) Under the model of IQ-tree automatic selection (automatic option in IQ-Tree) (Lam-Tung et al. 2015), The maximum likelihood phylogenetic TREE was deduced for 20000 ultra-fast guidance and approximate likelihood ratio tests using IQ-tree (Gascuel 2010; Minh et al. 2013), (5) For the construction of laccase family sequence of gray strain and white strain of Mushroom, phylogenetic tree was constructed by Bayesian method, and MrBayes was used under Wag+I+G model 3.2.6 Software reconstruction of Bayesian phylogenetic tree, sampling every 100 generations, discarding 25% of aging samples, remaining samples tree construction and calculation of posterior probability (Ronquist et al. 2012).

### 2.7 Analysis of the expression profiles of *HmLac*s in *H. marmoreus* based on RNA-seq

Total RNA was extracted from each sample according to the Kit (E.Z.N.A Plant RNA Kit, Omega, Biotech, Norcross, Ga). Illumina NEBNext® UltraTM RNA Library Prep Kit was used for Library construction, and the steps provided by the Kit were followed. Total RNA samples were commissioned to Be sequenced using Illumina HiSeqTM 2500 (Illumina Inc, CA, USA) platform by Beijing Nuohe Zhiyuan Bioinformatics Technology Co., LTD. Sequencing depth of each sample was 60X. At the same time, some RNA samples were kept in the −80°C refrigerator for storage.

The transcriptome data analysis steps are as follows: (1) all Illumina data sequenced by rna-seq were subjected to quality control by FastQC and Trimomatic (Bolger et al. 2014). (2) The HISAT2 (Kim et al. 2019) software was used to compare RNA sequencing read segments with default parameters, and Samtools (Li et al. 2009) was used to sort alignments according to the order of reference sequences, so as to obtain THE ALIGNMENT results in BAM format. Statistics were made according to the alignment results. (3) Based on the comparison results, Stringtie (Pertea et al.2015) software was used to estimate gene expression. In addition, Ballgown (Frazee et al. 2015) software provided by Stringtie were used to extract the number of read segments (read count) and FPKM values of genes located in gene exons after comparison from the results generated by Stringtie. (4) edgeR (Robinson et al. 2010) Read Counts was used to convert it into CPM (counts -per million) to filter genes that were not expressed or were all low expressed in all samples; (5) Differential expression analysis was conducted by Pairwise comparison of each sample using Bioconductor’s R language package edgeR: After filtering with CPM values, and Normalization with TMM (Mean of M-values), crosstalk significance is checked using statistical methods from edgeR (p values are calculated), The fold change between the two groups was estimated. Visualization of differentially expressed genes (DEG) using R language; (6) According to the list of differentially expressed genes, we used R language Bioconductor package topGO for GO enrichment analysis. TopGO estimates the significance of functional enrichment (p-value) based on the hypergeometric distribution into Fisher’s exact probability test. In the enrichment analysis results, p-value ≤0.05 was taken as the threshold, and the functions meeting this condition were defined as significantly enriched functions. (7) KEGG enrichment analysis was performed by R/Bioconductor package according to the list of differentially expressed genes. Enricher function was used to estimate the significance of functional enrichment (P value). With p-value ≤ 0.05 as the threshold, functions meeting this condition were defined as significantly enriched functions.

### 2.8 Experimental validation of *HmLacs* gene expression levels by qRT-PCR

The expression level of DEGs were validated by qRT-PCR. Primer3Plus (http://www.primer3plus.com/cgi-bin/dev/primer3plus.cgi) and NCBI Primer-BLAST (https://www.ncbi.nlm.nih.gov/tools/primer-blast/) were used to design gene-specific primer pairs (Table S1). Total RNA of tissues was extracted using Invitrogen Trizol, First-strand cDNA was synthesized with StarScriptIIFirst-strand cDNA Synthesis Mix with gDNA Remover for qPCR (A224-02; Genstar). The RT-qPCR was performed with 2x RealStar Green Fast Mixture (Genstar) on Multicolor Real-Time PCR Detection System (Bio-Rad). Reaction parameters for thermal cycling were 95°C for 2 min, followed by 40 cycles of 95°C for15 s and 60°C for 30 s, finally a melting curve (65-95°C, at the increments of 0.5°C) performed to confirm the PCR specificity. The expression level of each gene relative to housekeeping genes were calculated using the 2^-ΔΔCt^ with three replicates per sample (Li 2014). Then we made the correlated analyses between qRT-PCR values with FPKM values. In this experiment *GADPH* and *β-actin* were used as the reference genes.

## 3. Results and analysis

### 3.1 Identification of genes and characterization of homologous genes encode the laccase proteins in *H. marmoreus*

A total of 20 laccase genes were identified as candidates in the whole laccase genome by family alignment and similarity search against the published model fungi of *C. cinerea, P. ostreatus, F. velutipes*, and *L. edodes*. The comparison between CDD, SMRAT, and Swissport databases and analysis of laccase marker sequences (L1-L4) identified 18 laccase genes in the reference genome hm62-W of the white strain (Figure1); all these genes had three copper ion conserved domains. The amino acid sequences of 18 laccase genes were consistent with the characteristics of fungal laccase, and no deletion or replacement of amino acid residues was detected. Also, the characteristic sequence of fungal laccase was L1 – L4, i.e., the binding region of copper ions. At these sites, copper ions T1, T2, and T3 must bind to 10 histidines and 1 cysteine to form functional ligands (Figure 2).

**Figure 1.**
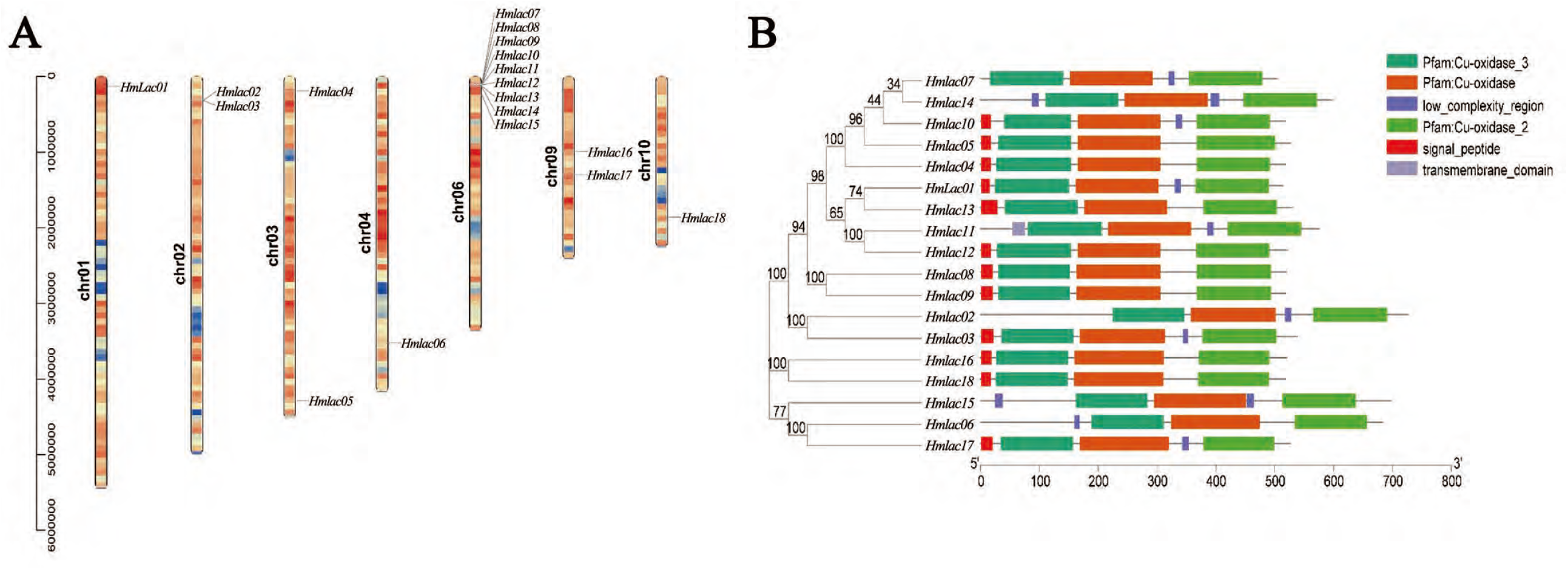
Distribution and domain of *HmLacs* on chromosomes. A. Chromosomal location and gene duplication of *HmLacs*. B. Phylogenetic tree and domain of *HmLacs*.

**Figure 2.**
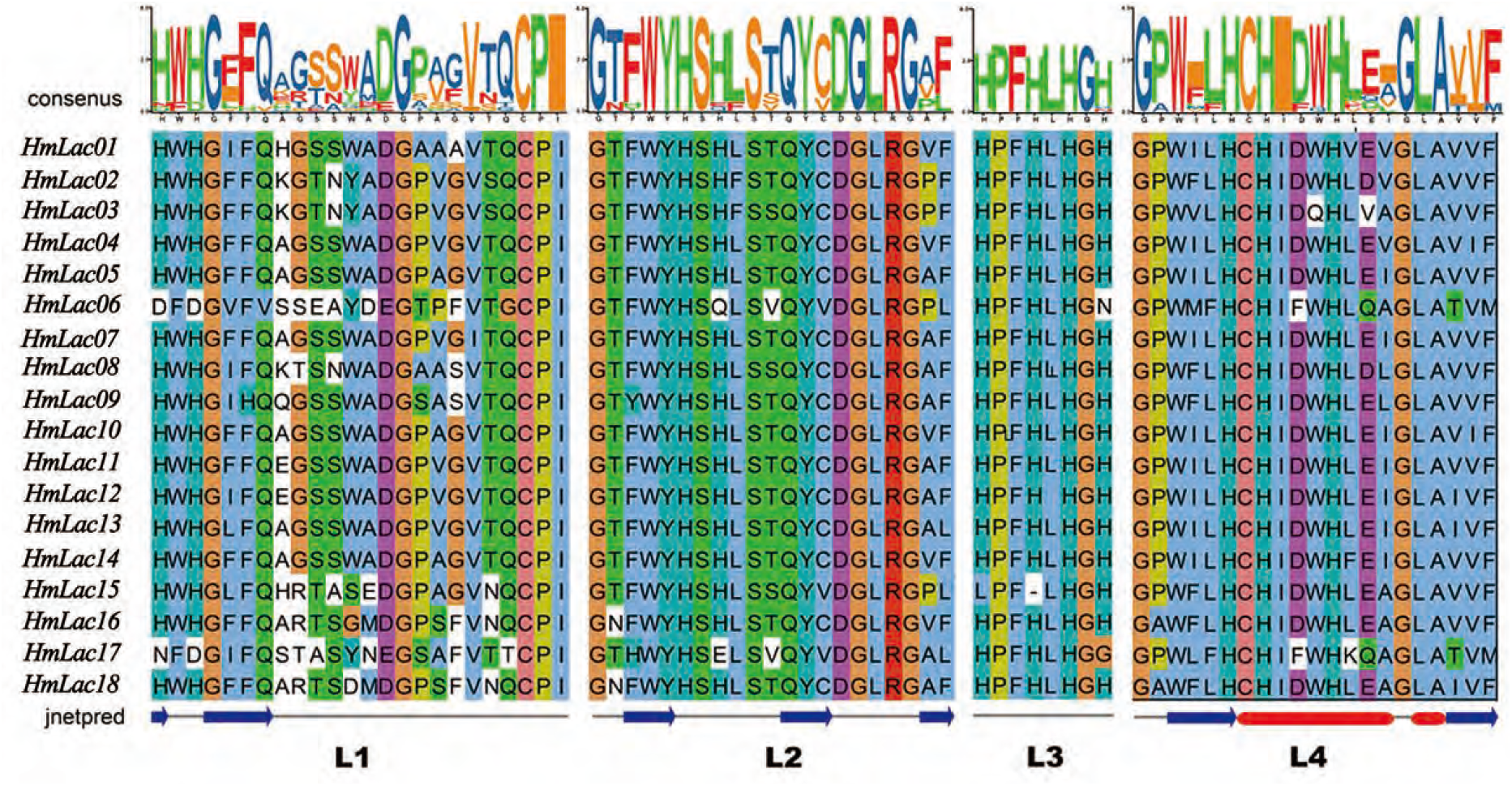
Standard sequence (L1-L4) multiple sequence alignment of *H. marmoreus* and Motif results.

The number of amino acids encoded by the 18 laccase gene proteins in the reference genome of laccase is 504 –726, and the molecular weight is 53.41356 – 80.99129 kDa (Table 1). The isoelectric point (IEP) was 4.41 – 6.52 except for *HmLac02*, which was 8.45. Compared to most fungal laccase, the IEP was in line with the physicochemical properties of fungal laccase. *HmLac02* was identified as a basic protein, which might have other functions of laccase.

**Table 1.** Features of *HmLacs* genes identified in *Hypsizygus marmoreus*

| Gene Name | Gene ID | Size( aa ) | MW |  | Chromosome No. | Signal Peptide | Intron/exon number |
| --- | --- | --- | --- | --- | --- | --- | --- |
|  |  |  | ( Da) | pI |  |  |  |
| *HmLac01* | *HM01gene000570* | 513 | 55329.78 | 6.2 | 1 | 16-17 | 15/14 |
| *HmLac02* | *HM01gene018960* | 726 | 80991.29 | 8.54 | 2 | - | 24/24 |
| *HmLac03* | *HM01gene018970* | 538 | 58712.2 | 5.52 | 2 | 22-23 | 14/14 |
| *HmLac04* | *HM01gene035380* | 518 | 55068.24 | 4.41 | 3 | 18-19 | 15/17 |
| *HmLac05* | *HM01gene051280* | 526 | 56023.63 | 4.99 | 3 | 18-20 | 15/20 |
| *HmLac06* | *HM01gene064720* | 683 | 74349.71 | 6.52 | 4 | - | 22/25 |
| *HmLac07* | *HM01gene082910* | 504 | 53413.56 | 4.45 | 7 | - | 15/21 |
| *HmLac08* | *HM01gene082960* | 520 | 56522.73 | 6.07 | 7 | 21-22 | 12/12 |
| *HmLac09* | *HM01gene082970* | 519 | 56568.31 | 3.4 | 7 | 21-22 | 12/16 |
| *HmLac10* | *HM01gene083010* | 517 | 54974.59 | 4.82 | 7 | 18-19 | 15/19 |
| *HmLac11* | *HM01gene083030* | 575 | 61995.57 | 3.41 | 7 | - | 18/18 |
| *HmLac12* | *HM01gene083050* | 521 | 55900.59 | 6.1 | 7 | 18-19 | 15/17 |
| *HmLac13* | *HM01gene083150* | 529 | 56965.73 | 5.94 | 7 | 29-30 | 15/21 |
| *HmLac14* | *HM01gene083210* | 597 | 63908.77 | 5.08 | 7 | - | 18/18 |
| *HmLac15* | *HM01gene083660* | 697 | 76917.7 | 6.05 | 7 | - | 28/28 |
| *HmLac16* | *HM01gene117180* | 520 | 57214.98 | 6.34 | 9 | 19-20 | 30/30 |
| *HmLac17* | *HM01gene118350* | 525 | 56864.6 | 6.56 | 9 | 21-22 | 22/24 |
| *HmLac18* | *HM01gene128560* | 519 | 56701.19 | 5.98 | 10 | 18-19 | 23/23 |

**Table 2.** List of the putative Motifs of *H. marmoreus HmLacs*.

The cleavage location of the laccase gene signal peptide was predicted, and the results showed that *HmLac01, HmLac03, HmLac04, HmLac05, HmLac08, HmLac09, HmLac10, HmLac12, HmLac13, HmLac16, HmLac17*. Twelve genes, including *HmLac18*, had signaling peptides at the *N*-terminal about 16 – 30 aa long, identified as secretory proteins. On the other hand, 6 kinase genes, *HmLac2, HmLac6, HmLac7, HmLac11, HmLac14*, and *HmLac15* did not harbor the typical signal peptides (Table 1). However, in the prediction of subcellular localization, the results showed that all 18 *HmLacs* were extracellular proteins, and the prediction of atypical laccase showed that they had atypical signaling peptides, presumably because not all members of the laccase gene family had the function of lignin degradation.

### 3.2 Genomic location and duplication events among *HmLac* genes

In the reference genome, 18 laccase genes were mainly distributed on chromosomes 1 (*HmLac01*), 2 (*HmLac02*), 3 (*HmLac04*), 5 (*HmLac05*), 4 (*HmLac06*), 6 (*HmLac07, HmLac*08, *HmLac09, HmLac10, HmLac11, HmLac12, HmLac13, HmLac14, and HmLac15*), chromosome 9 (*HmLac16* and *HmLac17*), and chromosome 10 (*HmLac18*). Interestingly, there are 9 *laccase* genes on chromosome 6 distributed in clusters, indicating that gene replication events occurred during the evolution of species. This phenomenon has not been reported previously (Figure 1). Herein, we also found that these 18 laccase genes, except *HmLac16* and *HmLac17* distributed in the middle of chromosome 9, were mainly distributed at both ends of the chromosome, and the chromatin in this region was loose, which needs to be investigated further. Combined with the MCScanX operation and manual correction results, we found that *HmLac10* and *HmLac14, HmLac11*, and *HmLac13* in the white line were lineal homologous genes. *HmLac02, HmLac03, HmLac08, HmLac09, HmLac10, HmLac11*, and *HmLac12* genes formed the tandem repeats, and *HmLac01, HmLac04, HmLac05, HmLac06, HmLac13, HmLac14, HmLac1 5, HmLac16, HmLac17*, and *HmLac18* may be produced by retrotransposition.

### 3.3 Gene structure, conserved motifs, and evolutionary correlations with *Hmlacs*

The comparison of the homology revealed that the consistency of amino acid sequence among laccase genes had a high diversity, and the identity ranged from 41.5–90.54%, which might be related to the specificity of functional differentiation in the evolutionary process (Figure S1). Nonmetric multidimensional scale (NMDS) analysis of the phylogenetic branch length showed that the white strain was divided into three groups, with significant differences among the groups (Figure S2). The phylogenetic tree of laccase gene construction of the *HM62-W* strain showed that the genes were divided into three subfamilies: Group 1 consisted of 13 family members, Group 2 had 2 family members, and Group 3 contained 3 family members. The amino acid similarity among each subgroup was low, which could be because different subfamilies are involved in diverse functions.

Gene structure analysisshowed that the number of exons of the laccase gene ranged from 9–30 and that the structures in each subfamily were similar (Figure 3). In Group 1, *G_HmLac10, G_HmLac13, G_HmLac4, HmLac12*, and *HmLac4* had 15 exons. The number of introns in Group 3 increased significantly, especially in the copper ion binding region (Cu-oxidase, Cu-oxidase_2, Cu-oxidase_3), suggesting that the introns might underlie the gene structure diversity, and the members of each group may perform different functions. Further analysis showed that all *HmLacs* genes contain at least one intron in their respective copper ion domain, which may be the conserved intron of the laccase gene. The Motif-based sequence analysis tool was used to identify 25 conservative motifs (length 6–50 aa) of proteins (Table S2). Among these conservative motifs, Group 3 has a unique Motif 20, 21, and 22, indicating that it might be related to the specific function of this group. The comparison of 25 conserved motifs in the CDD database revealed that 13 motifs had conserved domains of fungal laccase: Motif 1–6, 8, 10, 11, 12, 13, 23, and 24. These motifs were conserved across all *HmLacs*, while the rest were specific, providing favorable evidence for grouping *HmLacs*.

**Figure 3.**
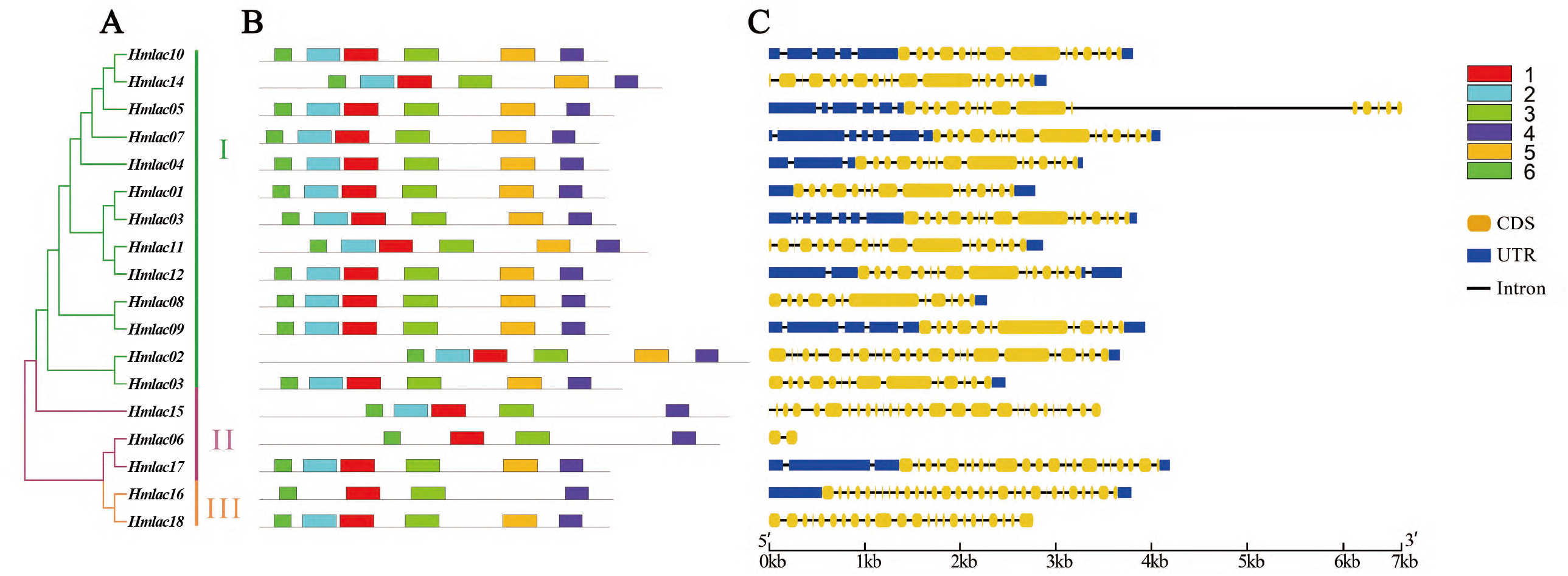
Phylogenetic tree, gene structure and conserved Motif analysis of 18 *HmLacs*. The phylogenetic tree was constructed based on *HmLacs* of *H. marmoreus* using Maximum Likelihood (ML) method, the branch labels designate bootstrap support values; b. The motif composition of *H. marmoreus HmLacs* proteins. Different colored boxes represent different motifs; c. Domain analysis of *H. marmoreus HmLacs* proteins.

### 3.4 Phylogenetic analysis and classification of *HmLacs*

To further categorize and investigate the evolutionary correlation of *HmLacs*, we identified 122 laccase domains from 64 *H. marmoreus* genomes which assembled by de novo using whole genome sequencing (WGS) data, and constructed an unrooted phylogenetic tree using the maximum likelihood (ML) methods. Based on the classification of *HmLacs* and the primary structural features of *HmLac* proteins, all 122 *HmLacs* genes were classified into three major groups and further divided into seven subgroups; the structure of each group of genes was similar (Figure 4).

**Figure 4.**
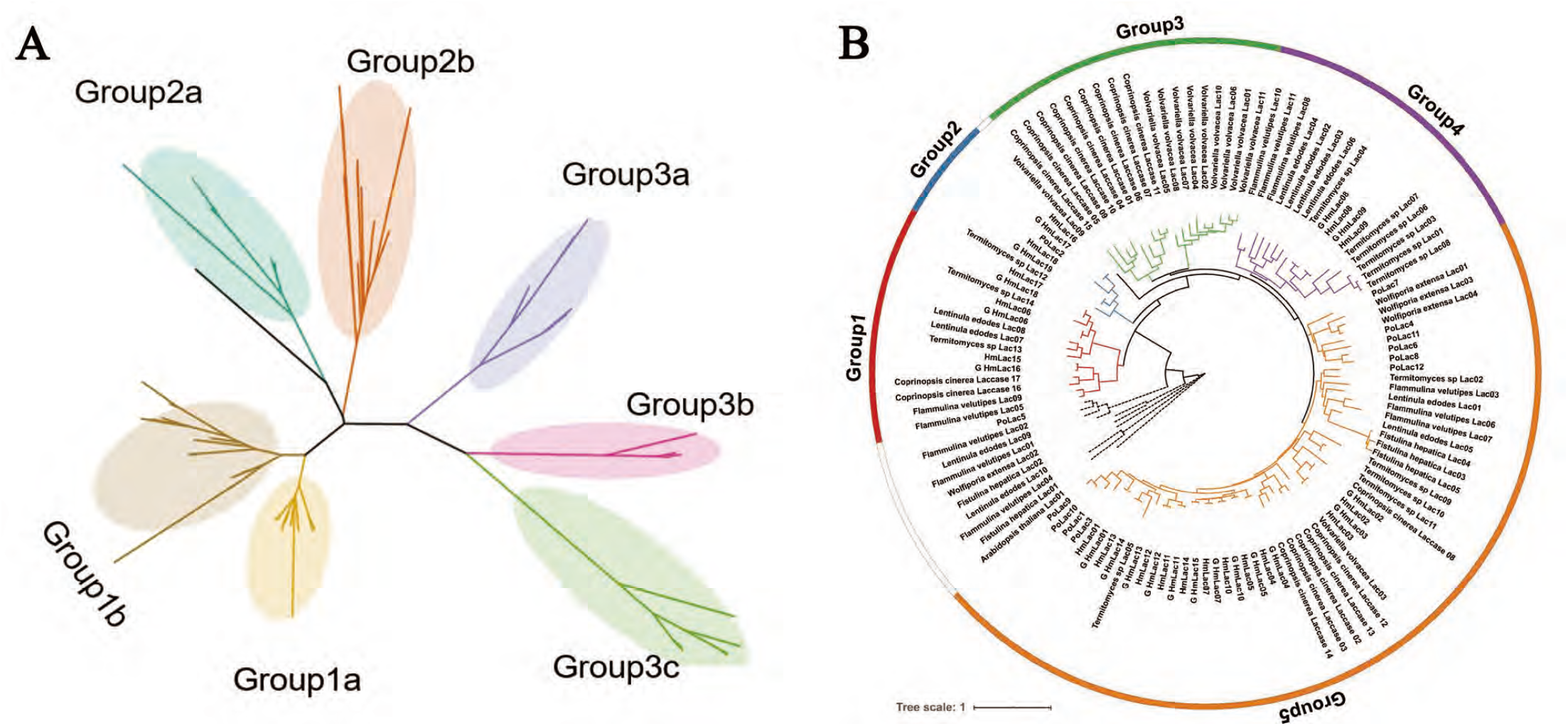
Phylogenetic tree using Maximum Likelihood (ML) analyses and class of *HmLacs* proteins. A. Phylogenetic tree of 122 laccase domains of 64 *H. marmoreus*. B. Phylogenetic tree of interspecific laccase gene family. The tree includes 10 species: *H. marmoreus, C. cinerea, M. albuminosa, P. ostreatus, L. edodes, F. velutipes, F. hepatica*, V. volvacea and *W. extensa*, and the Arabidopsis thaliana as outside groups.

The representative species of basidiomycetes, ligneous white-rot fungus (*C. cinerea, M. albuminosa, P. ostreatus, L. edodes*, and *F. velutipes*), ligneous brown rot fungus (*F. hepatica* and *W. extensa*), and rotting straw fungus (*V. volvacea*) were used to construct the phylogenetic trees, the *Arabidopsis thaliana* as outside groups (Figure 4). The results showed that the laccase genes are mainly divided into five groups, and each species’s laccase genes are clustered together respectively, indicating that there were gene replication events. Previous studies suggested that *PoLac2* was involved in lignin degradation and fruity body formation (Jiao et al. 2018). In Group 2, *PoLac2* was clustered with *HmLac16* and *HmLac18*, suggesting that the gene might be involved in lignin degradation. Group 3 did not consist of a laccase gene of *H. marmoreus*. The *HmLacs* genes are similar to the laccase genes in *M. albuminosa and P. ostreatus*, but unrelated to those of *F. hepatica, W. extensa*, and *V. volvacea*.

### 3.5 Expression profiles of *HmLac* genes in *H. marmoreus*

The life history of *H. marmoreus* can be divided into mycelium, mycelium kink (discoloring), primordial, bud, and forming stages. In the present study, the samples of the grey and white mycelium of *H. marmoreus* in the post-ripening (CK), kink (5 days after cap opening), and primordium formation stages (8 days after cap opening) were subjected to transcriptomic sequencing analysis.

Previous studies speculated that the white strain was the albino strain of the gray strain, which was weaker than the gray strain in growth speed and stress resistance. The expression levels of the three transcriptomes of *H. marmoreus* are shown in Figure 5. In the transcriptomic experiment, *HmLac09* and *HmLac10* were highly expressed in the gray strain, white mononuclear mycelia, and binuclear heterozygous mycelia, followed by *HmLac14*, suggesting that these genes are related to the growth and development of *H. marmoreus*. The expression levels of *HmLac02, HmLac05, HmLac08, and HmLac17* genes in gray or heterozygous strains were higher than those in the white mononuclear strains but did not differ significantly compared to the other laccase genes. In the cap epidermal transcriptome experiment, *HmLac09* gene was highly expressed in all three strains. The expression of *HmLac02, HmLac03, HmLac17, HmLac14*, and *HmLac18* in gray and hybrid was higher than that of the white strain, while the expression of *HmLac10* in white and hybrid was higher than that of the gray strain. The other 11 laccase genes were low expression, suggesting that they played a small role in the development of the cap. The results of transcriptome experiment III showed that *HmLac09* and *HmLac10* were highly expressed in different periods, further proving the critical role of these two genes in the growth and development of *H. marmoreus*. The expression levels of *HmLac02, HmLac05, HmLac08*, and *HmLac17* genes in gray or heterozygous strains were higher than those in white mononuclear strains and formed differential expression, which was consistent with the results of experiments I and II, indicating that any one of *HmLac02, HmLac05, HmLac08*, and *HmLac17* may be a factor leading to different growth rates of the gray strain in lignin decomposition. Taken together, *HmLac09, HmLac10, HmLac17*, and *HmLac18* are the key laccase genes in *H. marmoreus*.

**Figure 5.**
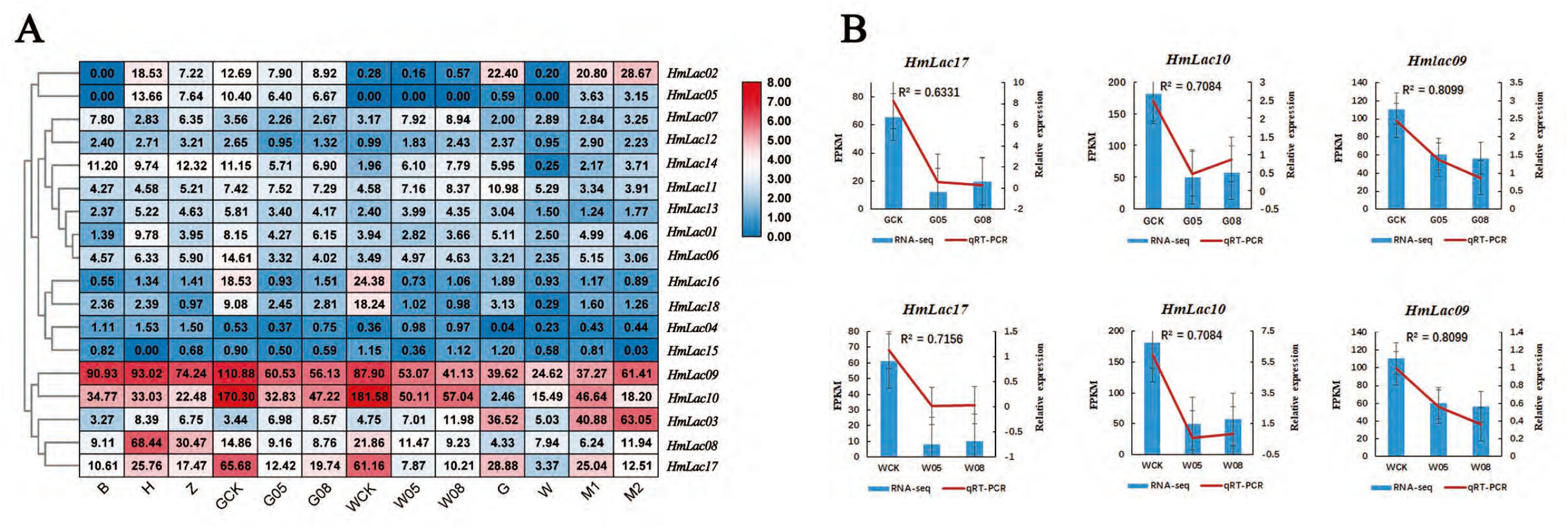
Expression patterns of 18 *HmLacs* genes expression profles and qRT-PCR analysis of the expression levels. A. Expression quantity heat map of 18 *HmLacs* in diffeent samples. B, H, and Z indicate Spores and dikaryon strain; CK, 05, 08 indicate post-ripening (CK), kink (5 days after cap opening), and primordium formation stages (8 days after cap opening) of *H. marmoreus* (white and gray strain); G and W, indicate white and gray strain, while M1 and M2 indicate crossing progenies.

The expression of *Lac17, Lac10*, and *Lac09* in different color strains was verified by Real-Time Quantitative Reverse Transcription (qRT-PCR) (Figure 5). These results showed that the correlation coefficients between Fragments Per Kilobase of exon model per Million mapped fragments (FPKM) and qRT-PCR were R^2^ = 0.7312(0.6331), R^2^ = 0.7084 (0.7084), and R^2^ = 0.8099 (0.8099) for *Lac17, Lac10*, and *Lac09*, respectively.

## 4. Discussion

Laccase of white-rot fungi is a major lignin-degrading gene that has been widely studied in recent years due to its high lignin-degrading efficiency. As one of the typical white-rot fungi, its nutritional components mainly originate from the degradation of lignin; hence, the laccase activity directly affects the production efficiency of *H. marmoreus* in the factory. Based on the whole genome identification, several laccase genes have been identified in many edible mushroom species, such as *P. ostreatus, F. velutipes* (Jiao et al. 2018; Wang et al. 2015). Based on the high-quality genome of *H. marmoreus*, this study compared the homologous sequences of related published species at the whole genome level, and combined with SMART database, CDD database, and laccase marker (L1-L4), identified 18 laccase family genes. This was 8 more than the 10 laccase genes identified by Zhang et al. through transcriptome assembly and splicing (Zhang et al. 2015). These 8 genes contain the characteristic structure of the copper ion binding region and are accurate members of the laccase gene family, which consists of 18 members. Currently, it is the largest laccase gene family found in Basidiomycota, exceeding the 17 laccase genes in *C. cinerea* (Kilaru et al. 2006), which might be the result of the growth and development of *H. marmoreus* and its dependence on laccase.

Based on the IEPs of the laccase gene family members, HmLac2 was identified as a basic protein. It is found that *H. marmoreus* often needs to grow in acidic environment, but some basic substances are secreted during the growth process, which gradually increases the pH value of the culture material. *HmLac2* plays a vital role in the special period. In addition, the mushroom has both acidic and basic proteins, indicating that its acid-base stability is different, which is consistent with the previous conclusion that fungal laccase has a wide optimal pH range (Baldrian 2006). Similar situations also occur in other species, such as *PoLac8* in *P. ostreatus*, thereby necessitating additional experiments are needed to verify the specific functions (Jiao et al. 2018). Some studies proposed that laccase was a secreted protein, but the results of signaling peptides in this experiment showed that some laccase did not have typical signal peptides, as in *F. velutipes, V. volvacea*, and *L. edodes* (Wang et al. 2015; Lu et al. 2015). Also, laccases might be intracellular enzymes with specific functions. Thus, we hypothesized that laccase, such as *HmLac2, HmLac6, HmLac7, HmLac11, HmLac14*, and *HmLac15* may be extracellular proteins without typical signaling peptides.

The distribution of 18 *HmLacs* on chromosome 6 was not uniform, and 9 genes were clustered linearly on chromosome 6, indicating that the laccase gene was derived from gene replication. The main driving force of the laccase gene family expansion was tandem repetition and reverse transcription transpose; 7 tandem repetition genes were identified. This phenomenon indicated that the original laccase genes were differentiated into paraphyletic homologs with different functions during the evolution process to meet the various functional requirements of fungi throughout the life cycle. Six lineal homologs have been identified in *H. marmoreus* and *P. ostreatus*, and it has been inferred that these genes may come from a common ancestor. The results of gene structure analysis showed that most of the *HmLacs* genes enriched in the same group had similar intron numbers. The results of gene structure analysis showed that most *HmLacs* genes enriched in the same group had similar intron numbers. The number and distribution of introns were related to gene evolution, which could be attributed to intron insertion or deletion caused by environmental pressure after species differentiation. In order to respond to various stresses promptly, genes must be activated quickly. In this respect, compact gene structures with fewer introns are conducive to expression (Kong et al. 2007).

The phylogenetic trees of laccase genes from 56 resequenced strains and 8 spore strains showed that the laccase gene family was divided into three large subgroups, which was consistent with a single reference genome laccase gene family. Among the species, 122 *HmLacs* were divided into five groups. In each group, the laccase genes were clustered together, indicating that the formation of laccase genes was earlier than speciation and that gene replication events occurred after species differentiation. The phylogenetic tree revealed that the laccase gene family of *H. marmoreus* is closely related to *L. edodes* and *F. velutipes*, and most *V. volvacea* and *C. cinerea* cluster into one branch, indicating that the laccases of *H. marmoreus* are closely related to wood rot fungi, but farther related to grass rot fungi, which is consistent with the previous findings.

*HmLac09* and *HmLac10* are highly expressed at various developmental stages and in the mononuclear mycelium and hybrid seed. *PoLac2* was overexpressed in *P. ostreatus* by *Agrobacterium*-mediated transformation (Jiao et al. 2018). The laccase activity of the transformant was increased to varying degrees, and the expression level of the *PoLac2* gene in the transformant was 2–8 times higher than that of the wild type. The lignin degradation rate of the transformant was 2.36–6.3% higher than that of the wild type within 30 days. In this study, the expression levels of *HmLac16, HmLac17*, and *HmLac18* reached their peak in the early bag-opening stage (post-ripening stage). Moreover, *HmLac16* and *HmLac17* were clustered on the same branch as *PoLac2*, which might be closely related to the lignin degradation of *H. marmoreus*.

## Author Contributions

Conceptualization, C.W., B.-B.L., F.L., and G.W.; methodology, Y.C.; formal analysis, Y.-Y.W., and H.-B.W.; investigation, B.-B.L.; data curation, Z.-J.S.; writing-original draft preparation, GW, B.-P.T., and L.M.; writing-review and editing, Y.C., C.W., and G.W. All authors have read and agreed to the published version of the manuscript.

## Institutional Review Board Statement

Not applicable.

## Informed Consent Statement

Not applicable.

## Data Availability Statement

The original genome data was uploaded to NCBI BioProject, under the accession number: PRJNA508399.

## Data availability

The genome sequences of H. marmoreus have been deposited at GeneBank under the accession number of JABWDO000000000. The data from this study were deposited with NCBI GenBank under accession numbers: PRJNA508399 and PRJNA644211

**Figure S1**. Consistency of amino acid sequences among *HmLacs*.

**Figure S2**. NMDS analysis of *HmLacs*.

**Table S1**. Laccase gene primer sequences.

